# Protein fusion strategies for a multi-component Rieske oxygenase

**DOI:** 10.1101/2024.06.09.598105

**Authors:** Michael E. Runda, Hui Miao, Sandy Schmidt

## Abstract

Rieske oxygenases (ROs) are enzyme systems involved in microbial biodegradation or late-stage modifications during natural product biosynthesis. A major obstacle to working with ROs is their dependence on multi-component electron transfer chains (ETCs). Thereby, electrons from NAD(P)H are shuttled directly via a reductase (Red) or indirectly via an additional ferredoxin (Fd) to a terminal oxygenase (Oxy) for oxygen activation and subsequent substrate conversion. The present work evaluates potential fusion strategies to simplify the ETC of the three-component cumene dioxygenase (CDO) from *Pseudomonas fluorescence*. In *in vitro* reactions, the fusion of CDO-Red to CDO-Fd is the most suitable for activation of CDO-Oxy with product formation of approximately 22 mM (72 % conversion). Furthermore, protein fusion to CDO-Oxy was found to be feasible, highlighting the versatility of the redox partner fusion approach. Overall, this study aims to contribute to the research field of ROs by providing a promising strategy to simplify their multi-component nature.

## INTRODUCTION

The class of Rieske oxygenases (ROs) comprises enzyme candidates capable of catalyzing various oxidation reactions using molecular O_2_.^1^ In nature, their reactivities are prominently involved in the biodegradation of aromatic xenobiotics in soil bacteria. Thereby, ROs facilitate the initial activation of arene substrates by catalyzing the formation of vicinal *cis*-dihydro diols.^2^ These compounds serve as intermediates, which, further processed by an enzymatic cascade, provide metabolites for the TCA cycle and thus contribute to the cell’s energy household.^3–5^

Besides their involvement in microbial biodegradation, more recent studies link ROs to natural product biosynthetic pathways.^6,7^ Thereby, ROs are proposed to perform late-stage functionalizations by catalyzing desaturations,^8^ cyclizations,^9,10^ dealkylations of heteroatoms,^11^ and hydroxylation reactions.^12^

This extensive repertoire of reactivities is based on the activation of O_2_ via single electrons from NAD(P)H.^13^ The supply of electrons to the active site of ROs is maintained by electron transfer chains (ETCs).^1^ These ETCs are composed of associated protein components, each harboring redox-active functional groups such as [2Fe-2S] clusters and/or flavins capable of accepting or donating single electrons.^14^ In the case of ROs, ETCs are composed of two- or three protein components, which allow the generally applied distinction between two- or three-component ROs.^15,16^ In all ROs, flavin-dependent reductases (RO-Reds) initially accept two electrons as hydride anion (H^-^) by oxidizing NAD(P)H.^17,18^ These electrons are then either directly,^19^ or indirectly via a small soluble [2Fe-2S] ferredoxins (RO-Fds)^20–23^ transferred to a terminal oxygenase (RO-Oxy). The RO-Oxys, which determine the catalytic function of ROs, are multi-component enzyme complexes that can be found as homotrimers (α_3_),^24–26^ homohexamers (α_3_α_3_),^27^ or heterohexamers (α_3_β_3_ or α_3_α’_3_).^28–32^ The α-subunits in Oxys harbor the distinctive Rieske [2Fe-2S] cluster, and a non-heme Fe coordinated by a facial triad.^33^ Electrons shuttled from the Rieske cluster to the non-heme iron center are utilized to form high-valent Fe-oxo intermediates upon O_2_ activation. Although the actual oxidation states of mononuclear Fe throughout the catalytic cycle of O_2_ activation are not fully understood, the high-valent oxo-iron intermediate is reported to be required for substrate oxidation. Due to the mode of O_2_ activation driven by the external supply of electrons via a redox chain, ROs are often associated with the well-studied cytochrome P450 proteins (CYPs).^34^ The CYP superfamily represents one of the oldest and largest classes of proteins, with members spread over all biological kingdoms.^35^ These enzymes are reported to play pivotal roles in diverse metabolic pathways such as steroid hormone biosynthesis^36^ and the degradation of xenobiotics.^37^ The latter function is commonly associated with human drug metabolism^38,39^ or the resistance of plants towards herbicides.^40^ CYPs are heme b-containing proteins that incorporate an atom of O_2_ into a substrate accompanied by the formation of H_2_O using electrons from NAD(P)H.^41^ This reaction cycle enables CYPs to catalyze a wide range of functionalization reactions such as hydroxylation, dealkylation, epoxidation, dehalogenation or desaturation on a variety of endogenous and exogenous substrates.^42^

The study of CYPs has been greatly facilitated by the introduction of fusion proteins such as the self-sufficient CYP102A1 (P450_BM3_), a fatty acid hydroxylase from *Bacillus megaterium*.^43,44^ P450_BM3_ is a soluble fusion protein consisting of a heme domain and an FMN- and FAD-containing reductase domain.^45^ This arrangement allows the catalysis of oxygenation reactions using only P450_BM3_ in the presence of NADPH and O_2_ without the addition of other redox partners. Due to this simplified reaction system, P450_BM3_ has served as a model for extensive enzyme engineering studies, addressing structural and mechanistic questions and contributing to the overall understanding of the catalytic cycles of CYPs.^46,47^ Driven by the described applicability advantage, research has recognized the feasibility of covalently linking redox partner proteins of CYPs and the advantage in terms of electron transfer properties and catalytic performance.^48–50^ In addition, the use of redox partner fusion proteins has been proposed to increase the applicability of CYPs and to identify new enzyme candidates.

Here, fusion strategies of physiological and non-physiological redox partners of the multi-component RO cumene dioxygenase (CDO) from *Pseudomonas fluorescence* are reported. CDO is a three-component RO and the CDO-Oxy (α_3_β_3_) is dependent on external electron supply from NADH via a reductase (CDO-Red) and a ferredoxin (CDO-Fd).^51^ As a proof of concept, we generated fusion constructs between CDO-Fd and CDO-Red as well as the reductase of the two-component phthalate dioxygenase (PDR) using a flexible GGGGS (G4S) linker. The construct with CDO-Red at the N-terminus (Red-G4S-Fd) was superior to the construct with the reverse redox partners (Fd-G4S-Red) in terms of CDO-Oxy activation, resulting in a 10-fold higher product formation in a biotransformation using indene (**1**) as a substitute (Scheme 1), with a total yield of more than 20 mM product (72% conversion).

**Scheme 1.**
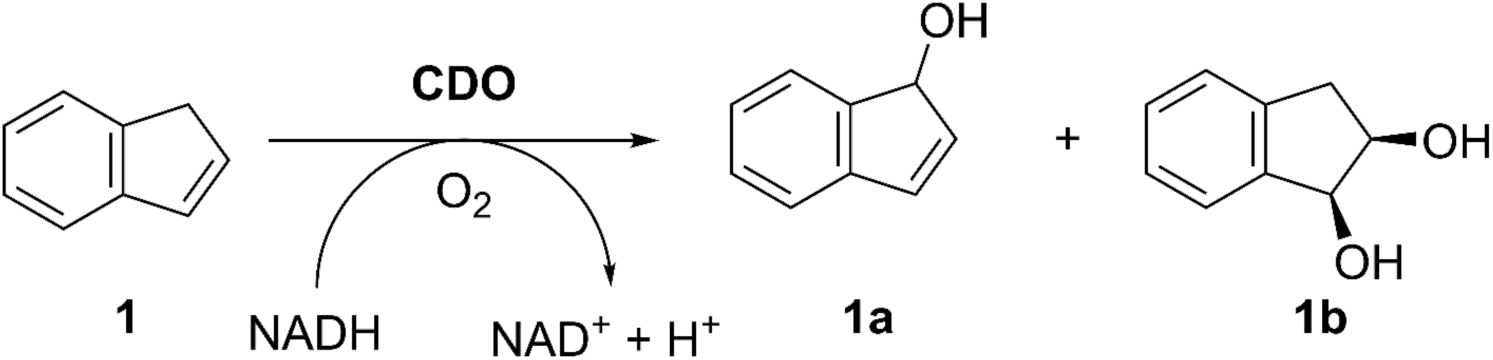
Conversion of indene (1) into 1*H*-indenol (1a) and *cis*-1,2,-indanediol (1b) catalyzed by CDO in the presence of molecular oxygen and NADH.

In addition to the highest conversion observed for CDO in a cell-free reaction system, the advantages in terms of total enzyme load are also highlighted. While an optimal redox partner ratio of the non-fused CDO system has been reported to be 1:7.5:2.5 (CDO-Oxy:CDO-Fd:CDO-Red),^51^ the *in vitro* system reported here showed the highest performance at equimolar concentrations between Red-G4S-Fd and CDO-Oxy. In addition, the feasibility of linking CDO-Red to the β-subunit of CDO-Oxy (Oxyβ-G4S-Red) further highlights the applicability of fusion strategies for multi-component ROs. To the best of our knowledge, this study is the first example of the implementation of artificial redox partner fusion constructs for the activation of RO-Oxys and represents a novel approach to improve the applicability of ROs.

## MATERIALS AND METHODS

### Protein Expression and Purification of RO components for *in vitro* biotransformation studies

Heterologous expression of (fused) RO protein components was performed using *E. coli* JM109(DE3) as the expression strain (Table S1). The genes encoding fusion protein constructs applied throughout this study were cloned in-frame to an N-terminal 6x His tag of a pET-28a(+) vector backbone (Table S2).

Initially, high-cell density precultures were prepared from cryo-glycerol stocks in baffled 250 mL Erlenmeyer flasks containing 50 mL LB supplemented with 50 µg mL^-1^ kanamycin. After approximately 16 h incubation at 37 °C and 200 rpm, the obtained high-cell density precultures were used to inoculate the main cultures at an OD_600_ of 0.1 for protein expression. The main cultures were performed in 1 L TB in 3 L baffled Erlenmeyer flasks, containing 50 µg mL^-1^ kanamycin. Protein expression was induced by adding 50 μM of β-D-1-thiogalactopyranoside (IPTG) at an OD_600_ of 1. After induction, the incubation temperature was lowered from 37 °C to 20 °C, maintaining shaking at 120 rpm for 18-20 h.

After cell harvesting at 4300 *g* and 4 °C for 30 min, cell pellets were washed by resuspending them in 100 mL lysis buffer (50 mM sodium phosphate buffer (SPB), 300 mM NaCl, 30 mM imidazole, 10 % glycerol, pH 7.2). After harvesting at 4300 *g* at 4 °C for 30 min and removing the supernatant, cell pellets were resuspended in 50 mL precooled lysis buffer.

Cell lysis was performed by sonication using a duty cycle of 50% and power control of 7 on the sonicator (Branson Sonifier 450). Sonication was performed in 8 cycles of pulsed sonication (15 seconds) separated by a 15-second pause. After cell lysis, centrifugation (18516 *g* at 4 °C) was performed for 1 h to obtain the cell-free extracts (CFEs) as supernatants.

CFEs were incubated with 2 mL (= 1 CV) Ni-sepharose for 1 hour at 4 °C. After removal of flow-through, each resin was washed with 5 CV of lysis buffer before eluting with elution buffer (50 mM SPB, 300 mM NaCl, 400 mM imidazole, 10% glycerol, pH 7.2). For non-fused CDO protein components, 0.5 to 2x CV elution fractions were collected for downstream sample preparation. For CDO fusion constructs, 0.5 to 3x CV elution fractions were collected during protein purification. Protein samples were desalted using desalting buffer (50 mM SPB, 300 mM NaCl, 10% glycerol, pH 7.2) on PD-10 desalting columns purchased from Cytiva (Danaher, Massachusetts, US).

Optional concentration of protein samples using Amicon^®^ Ultra centrifugal filters (Merck Millipore Ltd., Carrigtwohill, IRL) was performed prior to buffer exchange to reduce sample volume. Protein quantification was performed using a Coomassie-based colorimetric protein assay as described in a previous study.^51^ Protein samples were used directly in *in vitro* biotransformation reactions or stored at -20 °C after shock freezing in liquid nitrogen.

### Sample preparation for *in vitro* reactions

*In vitro* reactions were prepared in 20 mL tightly-sealed glass vials on a 1 mL scale. Unless otherwise stated, a standard reaction solution contained besides the CDO protein components, 1 mM of dithiothreitol (DTT), 1 mg mL^-1^ catalase from bovine liver (lyophilized powder, 2000-5000 units/mg protein, Sigma-Aldrich, Missouri, US) and 50 mM SPB (pH 7.2) as reaction buffer. To regenerate NADH, 10 U mL^-1^ glucose dehydrogenase (GDH) from *Bacillus megaterium* (expression and purification protocol described in the Supporting Information) were added in combination with 50 mM of D-glucose and 400 µM NAD^+^. The concentration of RO protein components and substrate applied in each reaction are indicated in the result section of the manuscript. In reactions exceeding 30 mM of **1**, 5 % DMSO was added as cosolvent. Reactions were performed in an incubation shaker at 30 °C and 120 rpm for 24 h. Reactions were stopped by the addition of solvent, extracted and product formation was quantified via GC-FID.

### Product quantification via Gas Chromatography (GC)

Products formed in *in vitro* reactions were quantified via GC-FID using a non-chiral OPTIMA™ 5MS column (Macherey-Nagel GmbH & Co. KG, Düren, Germany). Substrates and products were directly extracted from the aqueous reaction solutions with DCM containing 2 mM acetophenone as an internal standard. Reaction solutions were saturated with NaCl before liquid-liquid extraction to improve phase separation. The organic phase was dried over anhydrous MgSO_4_ before injecting into the GC-FID. The quantitative parameter applied for GC-FID can be found in the Supporting Information (Table S5).

## RESULTS AND DISCUSSION

To simplify the three-component CDO reaction system through a fusion strategy, we initially linked the CDO-Red to CDO-Fd covalently. The fusion constructs were designed with a flexible GGGGS linker (G4S) between the redox partners and an N-terminal 6x His-tag for downstream protein purification. To evaluate whether the purification tag or the linker affects the activity of one or both redox partners, we investigated both combinations, Fd-G4S-Red and Red-G4S-Fd (Scheme 2).

**Scheme 2.**
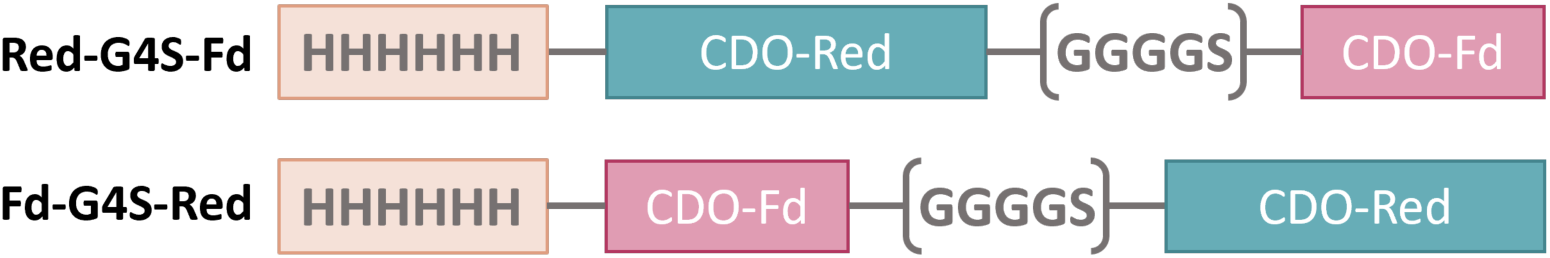
Design of fusion protein constructs (58.8 kDa) between the reductase (CDO-Red) and ferredoxin (CDO-Fd) of CDO. Domains are visualized in 5’→3’ direction. For purification purposes, the polyhistidine tag (6xHis) is attached to the N-termini and highlighted in orange. The flexible (G4S)-linker is indicated in brackets.

Both generated fusion constructs, Fd-G4S-Red and Red-G4S-Fd, were successfully expressed and purified (Figure S1 and S2), with a higher protein yield (∼ 1.5 fold) of the latter. Presumably, the well expressed CDO-Red favors the solubility of the downstream CDO-Fd, similar to solubility tagging strategies. While the elution fractions obtained during the purification of Fd-G4S-Red appeared yellow (Figure S1b), the analogous fractions containing purified Red-G4S-Fd showed an orange to brown color (Figure S2b). This color of the protein fractions is consistent with our previous observations from the isolation of other RO-Reds that contain a flavin domain as well as a [2Fe-2S] cluster.^52^

The ability of the fusion constructs to activate CDO-Oxy was evaluated in *in vitro* reactions using indene (**1**) as substrate (Scheme 1), which is converted to 1*H*-indenol (**1a**) and *cis*-1,2,-indanediol (**1b**). According to our hypothesis, the fusion constructs could bypass the need for Fd and Red as separate reaction components and thus directly mediate electrons from NADH to CDO-Oxy. Indeed, we observed product formation after 24 h, confirming that the fusion constructs mediate electron transfer to CDO-Oxy (Figure 1, left). Reactions with Red-G4S-Fd as the redox partner for CDO-Oxy yielded approximately 7.4 mM of product, 10-fold more than the fusion Fd-G4S-Red analog. As with *in vitro* reactions performed with CDO-WT,^51,52^ the presence of catalase as an hydrogen peroxide (H_2_O_2_) scavenger is essential to yield high product concentrations. Accordingly, only low substrate conversions of up to 1 % were observed in reactions without the addition of catalase (Figure 1, left).

**Figure 1.**
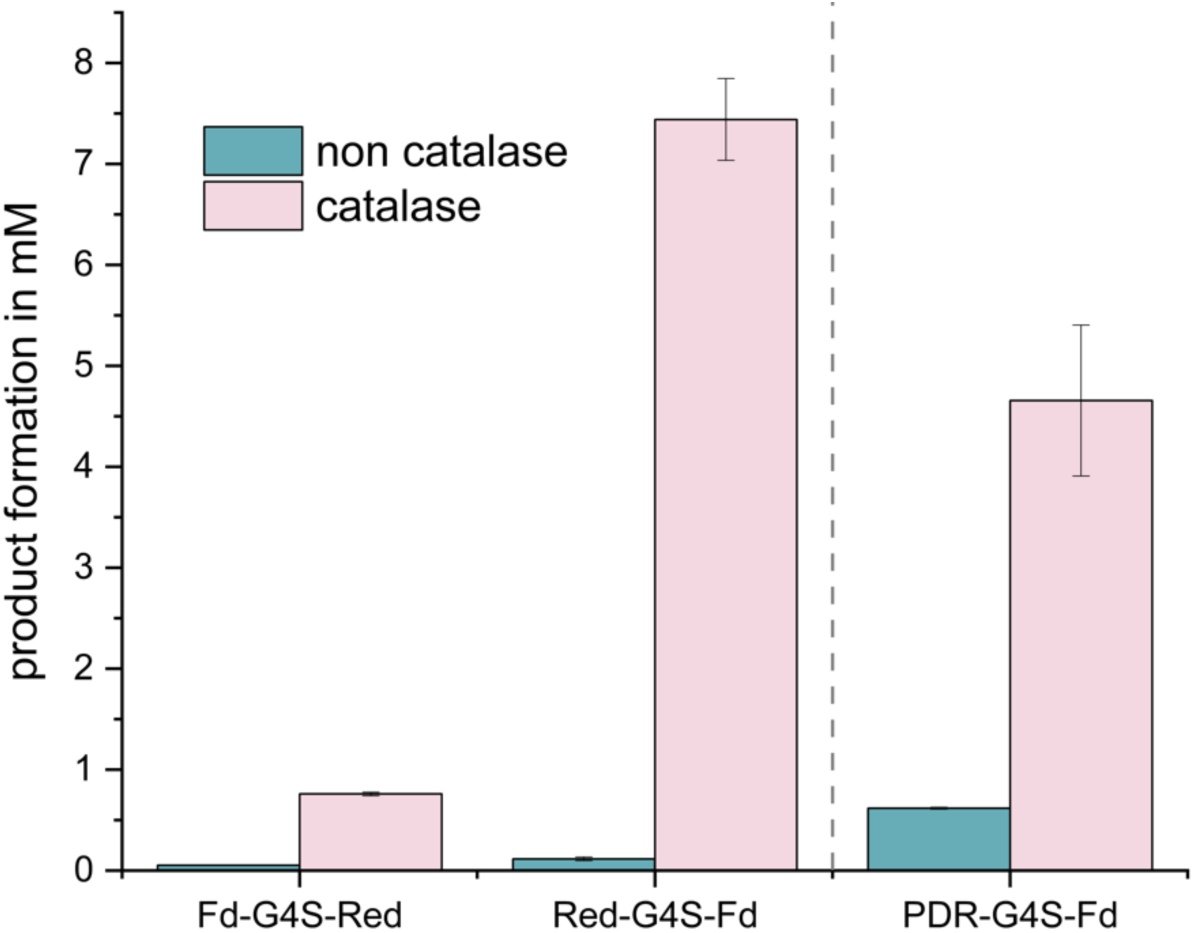
Product formations detected after 24 h in *in vitro* reactions using fusion proteins as redox partners for CDO-Oxy. Purified fusion proteins and CDO-Oxy were applied in equimolar amounts with concentrations of 4 μM each. *In vitro* reactions were performed in the presence or absence of 1 mg mL^-1^ catalase (see legend). Reaction conditions: 400 μM NAD^+^; 10 U mL^-1^ GDH, 50 mM glucose, 1 mM DTT, 10 mM (**1**), 50 mM SPB (pH 7.2) as reaction buffer. Incubation at 120 rpm and 30 °C for 24 h. Product formation corresponds to the sum of (**1a**) and (**1b**) determined via GC-FID (see Supporting Information for parameters).

The difference in expression levels and activity suggests that the arrangement of the two enzyme domains is critical, favoring the flavin domain of CDO-Red at the N-terminus. The superior Red-G4S-Fd fusion protein resembles the reductase of phthalate dioxygenase (PDR) at the primary sequence level with respect to the positions of the flavin and [2Fe-2S] prosthetic groups. PDR is a natural fusion harboring a flavin domain, a [2Fe-2S] domain and a NAD(P)H binding domain in close proximity.^17^ The arrangement of all domains within the protein backbon allows rapid electron transfer between the redox active centers. Furthermore, predicted 3D models of Fd-G4S-Red and Red-G4S-Fd (Figure 2) revealed significant differences in the spatial distances between the associated redox centers within both fusion constructs. Distances of 40.9 Å and 13.5 Å were determined in Fd-G4S-Red and Red-G4S-Fd, respectively. The latter distance is within the range (below 14 Å) where direct electron exchange is feasible^53^ and thus supports the hypothesis of rapid intramolecular electron transfer within Red-G4S-Fd.

**Figure 2.**
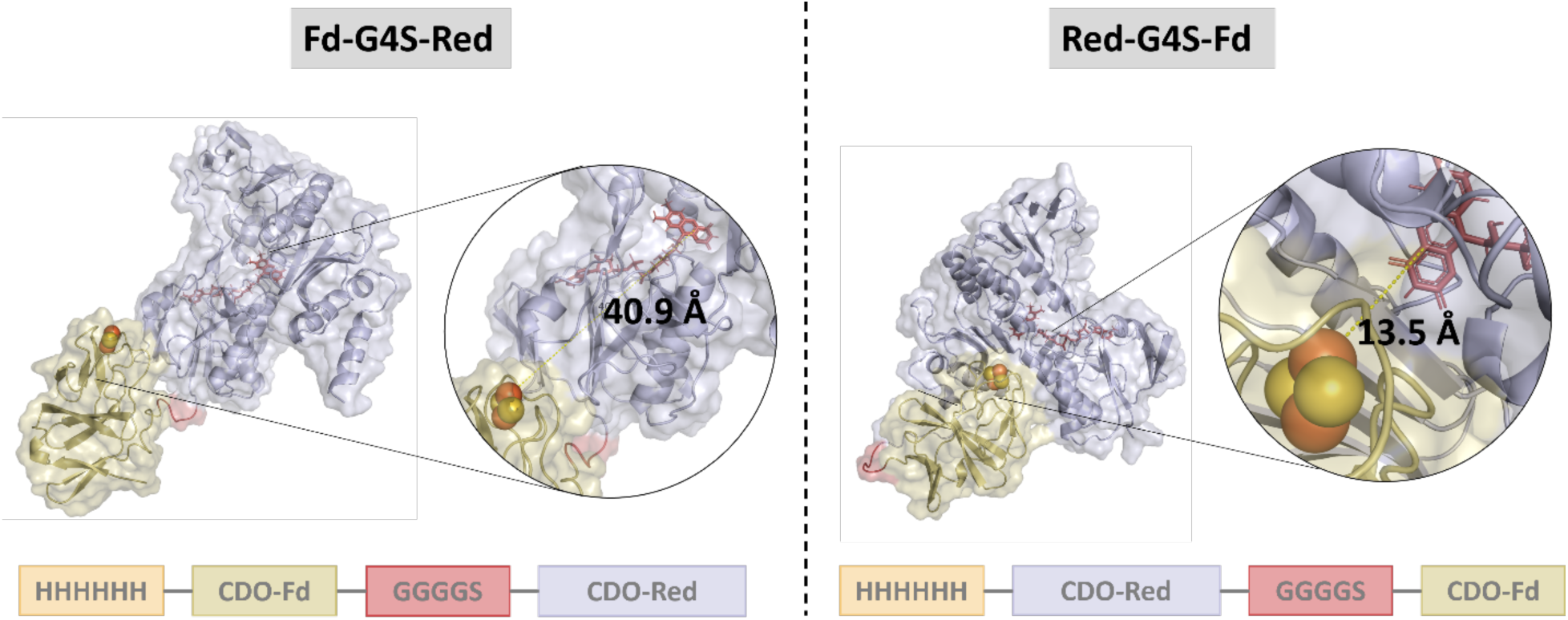
Predicted quaternary structures of Fd-G4S-Red (left) and Red-G4S-Fd (right). The spatial arrangements between the flavin prosthetic groups (shown as red sticks) and the associated [2Fe-2S] clusters (shown as spheres) are highlighted. The distances between the redox centers were measured to be 40.9 Å and 13.5 Å within the Fd-G4S-Red and Red-G4S-Fd protein scaffolds, respectively. For clarity, the arrangement of the protein subunits at the primary sequence level for each fusion construct is shown below, highlighted in the appropriate color. 3D structures were predicted using AlphaFold 3^54–56^ in combination with AlphaFill.^57^ Final structure refinement was performed using YASARA.

In a previous study, we emphasized the use of PDR in the context of CDO hybrid systems to enhance product formation in *in vitro* reactions.^52^ Based on these findings, we created a PDR-G4S-Fd protein fusion construct by exchanging the N-terminal CDO-Red domain of Red-G4S-Fd with PDR. Improved product formation (0.6 mM) was observed in the absence of catalase compared to reactions with the other fusion constructs, which is consistent with our previous findings with CDO hybrid systems (Figure 1, right).^52^ It is assumed that the rapid interdomain electron transfer within PDR suppresses O_2_ uncoupling and subsequent H_2_O_2_ formation. This results in a lower oxidative pressure and thus contributes to the stability of the *in vitro* RO system. On the other hand, reactions with PDR-G4S-Fd in the presence of catalase result in lower product formation compared to the CDO fusion Red-G4S-Fd (4.6 mM), suggesting that PDR is more affected by modifications at its N- or C-terminus than CDO-Red.

In addition to providing proof of concept, we further explored the advantages of fusion constructs over non-fused Fd and Red in an *in vitro* system for CDO. A clear advantage of using fused over non-fused redox components becomes apparent when evaluating the optimal protein ratio for bioconversion. The concentrations of CDO components to each other are critical and significantly determine the overall efficiency of *in vitro* reactions. Therefore, we performed reactions with varying concentrations of the fusion protein Red-G4S-Fd as redox partner for CDO-Oxy (Figure 3).

**Figure 3.**
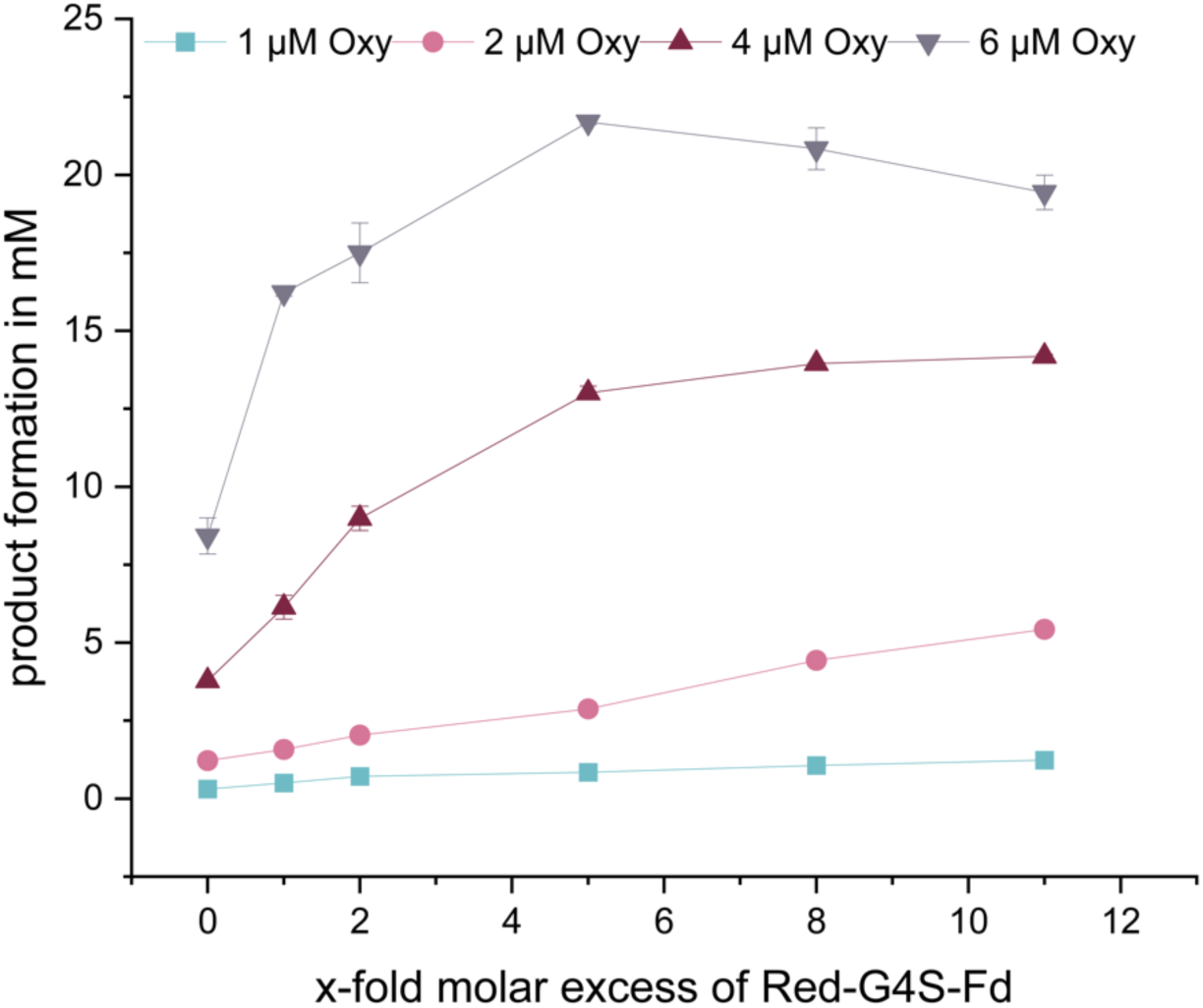
Product formation determined after 24 h in *in vitro* reactions using fusion redox partners and CDO-Oxy to evaluate the optimal protein ratio. Purified CDO-Oxy was applied in concentrations ranging from 1 to 6 μM (see legend at the top). The corresponding concentrations of Red-G4S-Fd are indicated as x-fold access over the concentration of CDO-Oxy. *In vitro* reactions were performed with 1 mg mL^-1^ catalase. Reaction conditions: 400 μM NAD^+^; 10 U mL^-1^ GDH, 50 mM glucose, 1 mM DTT, 5 % DMSO, 30 mM **1**, 50 mM SPB (pH 7.2) as reaction buffer. Incubation at 120 rpm and 30 °C for 24 h. Product formation corresponds to the sum of (**1a**) and (**1b**) determined via GC-FID (see Supporting Information for parameters).

We observed that the product concentrations increased with the given concentration of CDO-Oxy within the experimental range of 1 to 6 μM (Figure 3). Since CDO-Oxy is the catalytic component in terms of O_2_ activation and substrate conversion, its effect on product formation was expected. In addition, CDO-Oxy is a multi-component protein complex composed of 3 α- and 3 β-subunits. With catalytic sites embedded in the interface of two large α-subunits, each CDO-Oxy complex contains three active sites. The steep increase in product formation obtained from reactions with a ratio of 1:1 to 1:3 between the CDO-Oxy and the Red-G4S-Fd indicates a gradual saturation of the active sites with electrons. We conclude that all three reaction centers are simultaneously active and that the non-natural fusion proteins do not interfere with the redox partner interactions. Reactions with 4 μM of CDO-Oxy show a flattened product formation trend in the presence of Red-G4S-Fd above concentrations of 24 μM (Figure 3). Product concentrations measured in these reactions ranged from 13 to 14 mM. Under these conditions, the electron supply is saturated, and an increase in redox partners will not further contribute to enhanced bioconversion. In addition, the CDO-WT with an optimized redox partner ratio of 4 μM : 30 μM : 10 μM (Oxy : Fd : Red) yields in a product formation of approximately 4.6 mM under the same reaction conditions as previously reported.^52^ The significantly higher product formation observed in reactions with 4 μM CDO-Oxy in the presence of the Red-G4S-Fd fusion suggests a more efficient electron exchange between redox partners. Overall, the highest substrate conversion was obtained in reactions with a 1:6 ratio between CDO-Oxy (6 μM) and Red-G4S-Fd (36 μM). Approximately 73% of (**1**) could be converted by CDO, corresponding to a product concentration of 22 mM.

Furthermore, we evaluated the robustness of the *in vitro* fusion approach by performing reactions in the presence of 60 mM (**1**) and different ratios between CDO-Oxy and Red-G4S-Fd. The product formation plotted against the concentration of Red-G4S-Fd results in a saturation curve indicating the accumulation of non-interacting redox partners (Figure 4).

**Figure 4.**
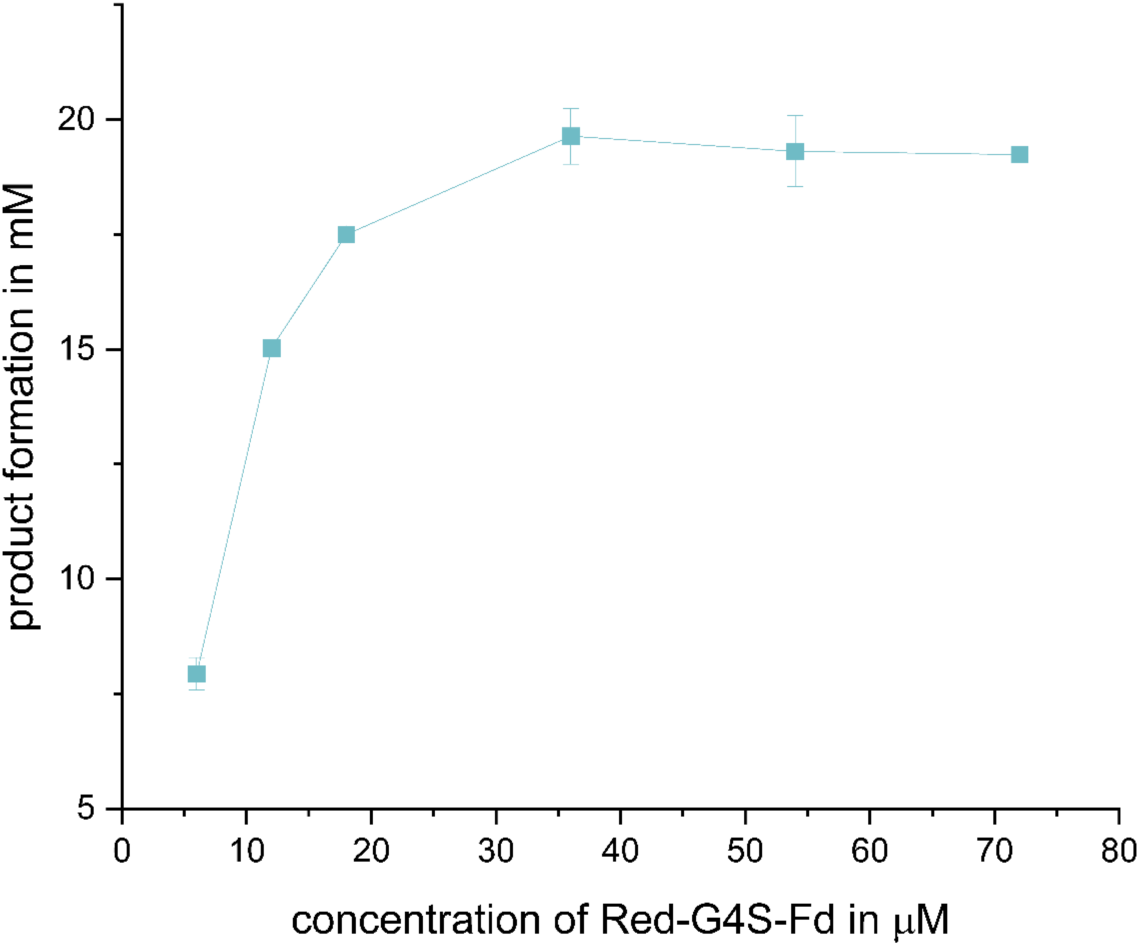
Product formations were determined after 24 h in *in vitro* reactions at elevated substrate concentration using fusion protein as redox partner for CDO-Oxy. Purified CDO-Oxy was applied in a concentration of 6 μM (see legend at the top). The corresponding concentrations of Red-G4S-Fd are indicated at the x-axis. *In vitro* reactions were performed with 1 mg mL^-1^ catalase. Reaction conditions: 400 μM NAD^+^; 10 U mL^-1^ GDH, 50 mM glucose, 1 mM DTT, 5 % DMSO, 60 mM **1**, 50 mM SPB (pH 7.2) as reaction buffer. Incubation at 120 rpm and 30 °C for 24 h. Product formation corresponds to the sum of **1a** and **1b** determined via GC-FID (see Supporting Information for parameters).

Using a substrate concentration of 60 mM (Figure 4), the product concentrations obtained are comparable to those obtained with 30 mM (**1**) (Figure 3), indicating that substrate inhibition on CDO can be ruled out. We believe that the total turnover number (TTN) of the enzyme is the limiting factor in maintaining the conversion of elevated substrate concentrations over time.

In addition to covalently linking the Fd and Red to form an active fusion construct, we further evaluated the CDO-Oxy as a potential site for fusion of redox partners. In contrast to the large α-subunit, which harbors the active site, the small β-subunit of α_3_β_3_-type RO-Oxys is thought to have primarily a structural function.^58,59^ In order not to influence the catalytic activity of CDO, we selected the β-subunit as a potential anchor site for the fusion of potential redox partners. Our previous observations suggest that the interaction of CDO-Fd with CDO-Oxy is more specific than its interaction with CDO-Red.^52^ Therefore, we decided to create the Oxyβ-G4S-Red construct, in which CDO-Red is covalently linked to the N-terminus of the β-subunit of CDO-Oxy via a G4S linker and the Fd is provided as free protein to ensure sufficient flexibility for electron transfer. The expression and purification of the Oxyβ-G4S-Red construct were judged to be successful based on the results of SDS-PAGE (Figure S4). To also evaluate the catalytic functionality of the Oxyβ-G4S-Red fusion protein, we performed *in vitro* reactions with 10 mM of (**1**). Reactions with 4 μM Oxyβ-G4S-Red in combination with 30 μM CDO-Fd yielded 2.8 mM product after 24 h in the presence of catalase. Although the observed product formation is significantly lower than with the Red-G4S-Fd approach (Figure 1), the information on CDO-Oxy as an anchor for potential fusion proteins is of high value for further optimization studies.

## CONCLUSION

To the best of our knowledge, this study represents the first example of successful protein fusion strategies for multi-component ROs. Covalent attachment of CDO-Fd to CDO-Red via a G4S linker results in functional fusion proteins capable of activating CDO-Oxy. In our study, the Red-G4S-Fd fusion construct, in which CDO-Fd is attached to the C-terminus of CDO-Red, is the more efficient construct for the conversion of (**1**). With more than 20 mM product formation (72% conversion) after 24 hours, the advantage of using Red-G4S-Fd is evident. In addition to the overall simplification of the CDO system, the use of Red-G4S-Fd further demonstrates its potential to reduce the total protein required for conversion. Compared to the unfused CDO, for which an optimal molar ratio of CDO-Oxy : CDO-Fd : CDO-Red of 1 : 7.5 : 2.5 was determined, a ratio of 1 : 6 between CDO-Oxy : Red-G4S-Fd resulted in the highest product formation in 24 h. Furthermore, we were able to successfully fuse the CDO-Red to the small subunit of CDO-Oxy (Oxyβ-G4S-Red). Despite minor product formation observed in *in vitro* reactions in the presence of CDO-Fd, it can serve as a valuable template for further optimization studies. Overall, the applicability of the fusion strategies described herein is expected to contribute significantly to the research field of ROs.

## Notes

The authors declare no competing financial interest.

## ABBREVIATIONS

CDO: cumene dioxygenase
DTT: dithiothreitol
Fd: ferredoxin
H_2_O_2_: hydrogen peroxide
IPTG: isopropyl ß-D-1-thiogalactopyranoside
LB: *lysogeny broth*
Oxy: terminal RO oxygenase
PDR: phthalate dioxygenase reductase, Red, reductase
RO: Rieske oxygenase
ROS: reactive oxygen species
SPB: sodium phosphate buffer
TB: terrific broth.

