## Supporting Information for "Protein fusion strategies for a multi-component Rieske oxygenase"

### List of Abbreviations

| | |
|---|---|
| CDO | cumene dioxygenase |
| c.o. | codon-optimized |
| DCM | dichloromethane |
| DMSO | dimethyl sulfoxide |
| GC | gas chromatography |
| Fd | ferredoxin |
| FID | flame ionization detector |
| GDH | glucose dehydrogenase |
| DMSO | horseradish peroxidase |
| IPA | integrated peak area |
| IST | internal standard |
| ORF | open reading frame |
| Red | Reductase |
| RO | Rieske oxygenase |
| RT | retention time |
| RRF | relative response factor |

### Materials

Unless stated otherwise, all chemicals and solvents were purchased from Sigma-Aldrich (St. Louis, USA), Merck KGaA (Darmstadt, Germany), TCI-Tokyo Chemical Industry (Tokyo, Japan), Carl Roth GmbH and Co. KG (Karlsruhe, Germany) and Duchefa Biochemie (Haarlem, The Netherlands) in the highest purity available. Enzymes for Gibson Assembly were purchased from New England Biolabs (Ipswich, Massachusetts, US). FastDigest™ enzymes from Thermo Scientific (Massachusetts, USA) were used for restriction digestion of DNA. Kits for PCR product purification, DNA gel extraction, and plasmid isolation were used from Qiagen or MACHEREY-NAGEL (Düren, Germany). The service of Macrogen Europe (Amsterdam, The Netherlands) was used for DNA sequencing. Synthetic genes were purchased from TWIST Bioscience (South San Francisco, California, US). GC-FID analyses were performed on a Shimadzu GC-2010 Plus equipped with an AOC-20i auto-injector. Column specifications and conditions for GC analyses are detailed in another section of the Supporting Information.

### Bacterial Strains

Bacterial strains used for cloning and protein purification throughout the project are summarized in **Table S1**.

**Table S1.** ***E. coli*** **strains used for molecular cloning and protein expression.**

| Strain | Genotype | Description/Use |
|---|---|---|
| **E. coli DH5α** | $F^{-}$ φ80lacZΔM15 Δ(lacZYA-argF)U169 recA1 endA1 hsdR17($r_K^{-}$, $m_K^{+}$) $gal^{-}$phoA supE44 $λ^{-}$thi-1 gyrA96 relA1 | routine cloning |
| **E. coli TOP10** | $F^{-}$*mcr*A Δ(*mrr-hsd*RMS-*mcr*BC) φ80*lac*ZΔM15 Δ*lac*X74 *rec*A1 *ara*D139 Δ(*ara-leu*)7697 *gal*U *gal*K $λ^{-}$ *rps*L($Str^{R}$) *end*A1 *nup*G | routine cloning |
| ***E. coli* JM109 (DE3)** | *end*A1, *rec*A1, *gyr*A96, *thi, hsd*R17 ($r_k^{-}$, $m_k^{+}$), *rel*A1,*sup*E44, λ–, Δ(*lac-pro*AB), [F´, *tra*D36, *pro*AB, *lac*$I^{q}$ZΔM15], lDE3 | expression host for Rieske oxygenase (RO) protein components |
| ***E. coli* BL21(DE3)** | $F^{-}$*omp*T *hsdS*$_B$ ($r_B^{-}$, $m_B^{-}$) *gal dcm* (DE3) | expression host for GDH |

### Plasmid Constructs

#### Plasmid constructs for protein expression

Plasmid constructs used for the heterologous expression of protein components are summarized in **Table S2**.

**Table S2. Plasmid constructs for heterologous expression in *E. coli*.**

| Construct | Protein expressed | Insert | Comment |
|---|---|---|---|
| **pET28a-[N-His]-CumA1_CumA2** | non-fused Oxy component of CDO | *cumA1* (N-terminal 6xHis-tag, c.o.), *cumA2* (c.o.) | Created previously[1] |
| **pET28a-[N-His]-CumA3** | non-fused Fd component of CDO | *cumA3* (N-terminal 6xHis-tag, c.o.) | Created previously[1] |
| **pET28a-[N-His]-CumA3** | non-fused Red component of CDO | *cumA4* (N-terminal 6xHis-tag, c.o.) | Created previously[1] |
| **pET28a-[N-His]-CumA3-CumA4** | Fd-G4S-Red fusion construct | *cumA3-G4S-cumA4* (N-terminal 6xHis-tag, c.o.) | Created by TWIST Biosience |
| **pET28a-[N-His]-CumA4-CumA3** | Red-G4S-Fd fusion construct | *cumA4-[G4S]-cumA3 (*N-terminal 6xHis-Tag, c.o. genes | Created by TWIST Biosience |
| **pET28a-[N-His]-PDR-CumA3** | PDR-G4S-Red fusion construct | *pdr-[G4S]-cumA3 (*N-terminal 6xHis-Tag, c.o. genes | Created by TWIST Biosience |
| **pET28a-[N-His]-CumA1_CumA4-CumA2** | $Oxy_\beta$-G4S-Red fusion construct | *cumA1* (N-terminal 6xHis-tag, c.o.), *cumA4-[G4S]-cumA2* (c.o.) | Created via Gibson assembly |
| **pKTS-GDH** | Glucose dehydrogenase (GDH) from *Bacillus megaterium* | *gdh (*N-terminal 6xHis-tag) | Provided by Prof. M. E. Brenna, see reference[2] |

#### Creating the fusion construct for the expression of $Oxy_\beta$-G4S-Red

To create the pET28a-[N-His]-CumA1_CumA4-CumA2 plasmid (**Table S2**) for the heterologous expression and purifcation of $Oxy_\beta$-G4S-Red fusion construct, Gibson assembly was performed. Therefore, DNA fragments for assembly have to be provided in a linear form with suitable Gibson overhangs. In the following, the preparation of the large backbone

**Fragment 1** (pET28a(+) backbone with *cumA1* and *cumA2*) and insert **Fragment 2** (harboring CumA4 with Gibson overhangs) is described.

**Preparation of Fragment 1 (7324 bp)**

**Fragment 1** was obtained by PCR amplification of pET28a-[N-His]-CumA1_CumA2 (**Table S2**). Therefore, primers P1 and P2 (**Table S3**) were designed which bind at the 5'-end of *cumA2* and the 3'-end of *cumA1*, respectively. The obtained PCR product represents the pET28a-[N-His]-CumA1_CumA2 template, linearized between *cumA1* and *cumA2*. To eliminate the template DNA, DpnI digestion was performed (37 °C for 1h) followed by gel purification. Isolated PCR product was directly used for Gibson assembly.

**Preparation of Fragment 2 (1233 bp)**

**Fragment 2** was obtained by two rounds of PCR amplification using pET28a-[N-His]-CumA4 (**Table S2**) as a template. In the first round, *cumA4* was amplified using primers P3 and P4. After gel purification, the obtained PCR product was applied as the template for the second round. Therefore, primers were designed (P5 and P6) binding at both ends of *cumA4* and creating suitable Gibson overhangs. After gel purification, **Fragment 2** was directly used for Gibson assembly.

**Gibson Assembly**

Gibson assembly was performed with equimolar amounts of **Fragment 1** and **Fragment 2**. Therefore, calculated amounts of DNA solutions (in total 10 µL) were added to a 15 µL aliquot of self-made Gibson master mix. After 1 h incubation at 50 °C, 5 µL of the Gibson reaction mixture was used in heat-shock transformation of competent *E. coli* TOP10 cells. Colonies containing the correct construct were selected by colony PCR followed by restriction analysis and finally sequencing of the whole plasmid.

Oligonucleotides applied in the PCR amplification for Gibson assembly are summarized in Table S3.

**Table S3. Oligonucleotides used in Gibson assembly**

| Name | Sequence 5'→ 3' | Description |
| --- | --- | --- |
| **P1** | GTGGTTCCACCAGCGCAGATCTGAC | Forward primer used to create **Fragment 1** for Gibson assembly |
| **P2** | GGTATATCTCCTTCTTAAAGTTAAAC | Reverse primer used to create **Fragment 1** for Gibson assembly |
| **P3** | ATGATTAAAAGCATCGTGATTATTGG | Forward primer used in PCR round 1 to create **Fragment 2** for Gibson assembly |
| **P4** | CACCACCCTCGCAACGTTCGGCTTTTG | Reverse primer used in PCR round 1 to create **Fragment 2** for Gibson assembly |
| **P5** | AATAATTTTGTTTAACTTTAAGAAGGAGATATACCATGATTAAAAGCATCGTGATTATTGGTG | Forward primer used in PCR round 2 to create **Fragment 2** for Gibson assembly |
| **P6** | AATCGGTTTGGTCAGATCTGCGCTGGTGGAACCACCACCACCCTCGCAACGTT | Reverse primer used in PCR round 2 to create **Fragment 2** for Gibson assembly |

### Protein Purification

#### Expression and Purification of Glucose dehydrogenase (GDH)

Heterologous expression of GDH from *Bacillus megaterium* was performed using *E. coli* BL21(DE3) as the expression strain. The pKTS-GDH plasmid containing GDH in frame with an N-terminal 6xHis-Tag was provided by Prof. M. E. Brenna (**Table S2**).

Precultures were prepared in baffled Erlenmeyer flasks by inoculating LB liquid media supplemented with 50 µg $mL^{-1}$ ampicillin with a small amount of bacterial glycerol stock. After approximately 16 h incubation at 37 °C at 200 rpm, the obtained high cell density cultures were used to inoculate the main cultures for protein expression an $OD_{600}$ of 0.1. The main cultures were performed in 1 L LB in 3 L baffled Erlenmeyer flasks, containing 50 µg mL-1 ampicillin. Protein expression was induced by adding 100 µM of β-D-1-thiogalactopyranoside (IPTG) at an $OD_{600}$ of 0.6 – 0.8. After induction, the incubation temperature was lowered from 37 °C to 20 °C, maintaining shaking at 120 rpm for 18-20 h.

After cell harvesting at 4300 *g* at 4 °C for 30 min, cell pellets were washed by resuspending them in 100 mL of lysis buffer (50 mM sodium phosphate buffer (SPB), 300 mM NaCl, 30 mM imidazole, 10 % glycerol, pH 7.2). After harvesting at 4300 *g* at 4 °C for 30 min and removing the supernatant, cell pellets were resuspended in 50 mL of precooled lysis buffer.

Cell lysis was performed by sonication using a duty cycle of 50% and power control of 7 on the sonicator (Branson Sonifier 450). Sonication was performed in 8 cycles of pulsed sonication (15 seconds) separated by a 15-second pause. After cell lysis, centrifugation (18516 *g* at 4 °C) was performed for 1 h to obtain the cell-free extracts (CFEs) as supernatants.

CFEs were incubated for 1 h with 2 mL (= 1CV) of Ni-sepharose at 5 °C. After removing the flowthrough, each resin was washed with 5 CV of lysis buffer before eluting with elution buffer (50 mM SPB, 300 mM NaCl, 400 mM imidazole, 10 % glycerol, pH 7.2). The elution fractions containing GDH protein (evaluation via SDS-PAGE, see Figure S5) were pooled and subsequently desalted using PD-10 desalting columns purchased from Cytiva (Danaher, Massachusetts, US) using a desalting buffer (50 mM SPB, 300 mM NaCl, 10 % glycerol, pH 7.2).

Optional concentration of protein samples with Amicon® Ultra centrifugal filters (Merck Millipore Ltd., Carrigtwohill, IRL) was performed before buffer exchange to lower sample volumes. The activity of GDH was determined measuring the regeneration of NADH at 340 nm in a plate reader.

Protein samples were directly used in *in vitro* biotransformation reactions or stored at -20 °C after shock-freezing in liquid nitrogen.

#### SDS-PAGE to evaluate protein expression and purification

Figures **S1** to **S4** show **the** stained SDS-PAGE gels of purified fusion proteins used throughout this study.

a)

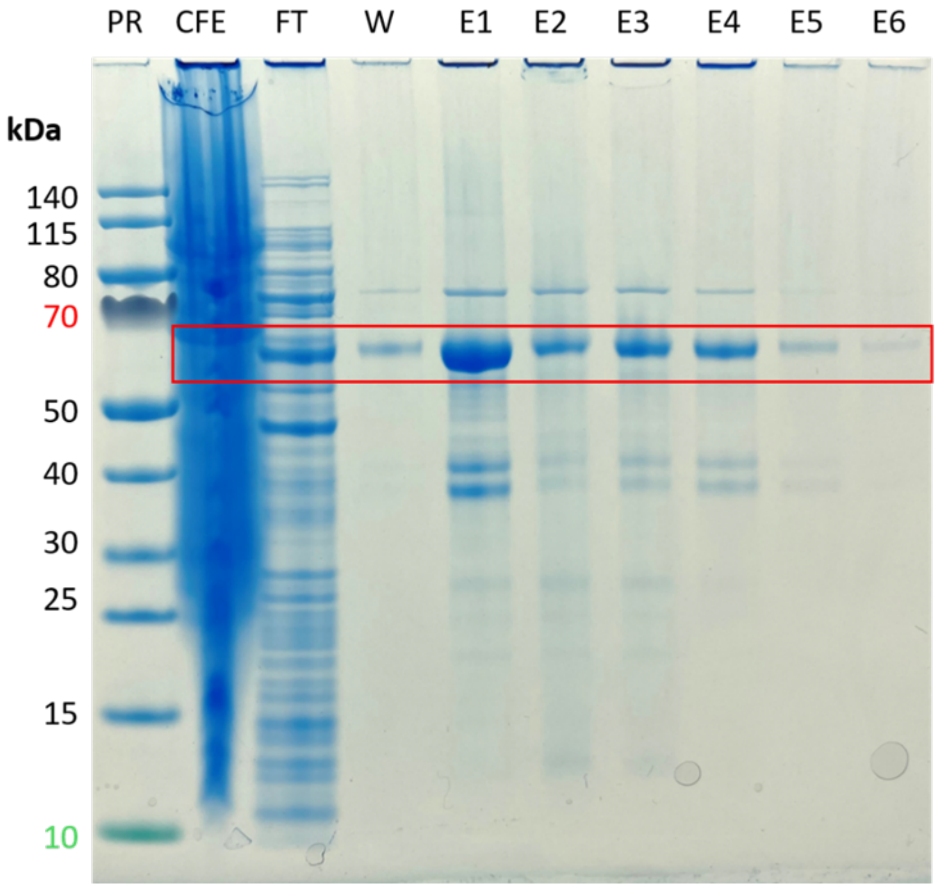

b)

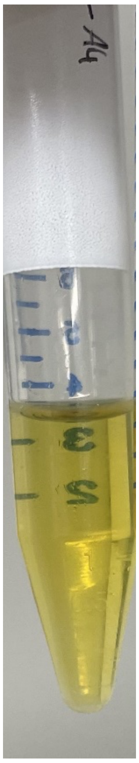

**Figure S1a. Purification of Fd-G4S-Red fusion protein construct (58.8 kDa).** a) Stained SDS-PAGE showing protein fractions obtained during the purification procedure of Fd-G4S-Red fusion protein construct (58.8 kDa). The expected height for bands representing the desired protein is highlighted in red. PR: PageRuler™ protein ladder, CFE: cell-free extract, FT: flow through, W: washing fraction, E1-E6: elution fractions (0.5 CV of elution buffer). NuPAGE™ Bis-Tris Mini Protein Gels, 4–12 %, Running conditions: 150 V for ~60 min in MES running buffer. b) Purified and desalted protein sample.

a)

b)

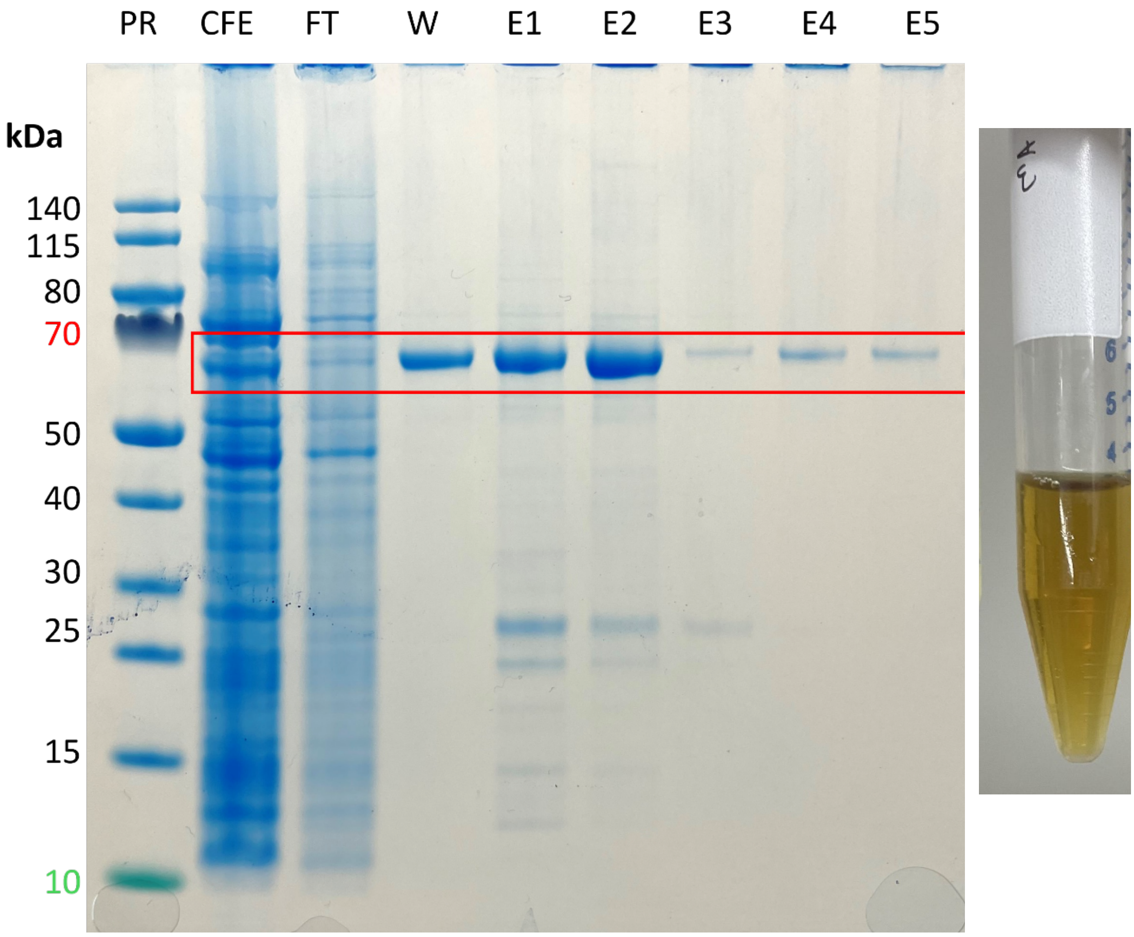

**Figure S2b. Purification of Red-G4S-Fd fusion protein construct (58.8 kDa).** Stained SDS-PAGE showing protein fractions obtained during the purification procedure of Red-G4S-Fd fusion protein construct (58.8 kDa). The expected height for bands representing the desired protein is highlighted in red. PR: PageRuler™ protein ladder, CFE: cell-free extract, FT: flow through, W: washing fraction, E1-E5: elution fractions (0.5 CV of elution buffer). NuPAGE™ Bis-Tris Mini Protein Gels, 4–12 %, Running conditions: 150 V for ~60 min in MES running buffer. b) Purified and desalted protein sample.

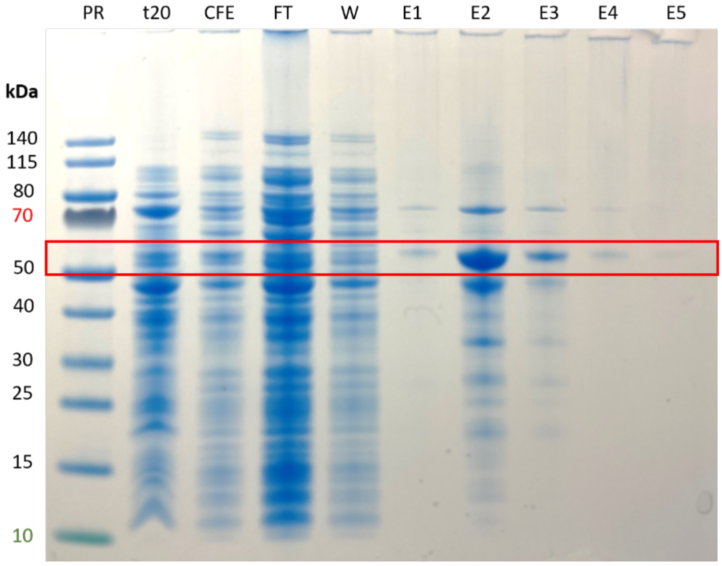

**Figure S3.** Stained SDS-PAGE showing protein fractions obtained during expression and the purification procedure of PDR-G4S-Fd fusion protein construct (51.2 kDa). The expected height for bands representing the desired protein is highlighted in red. PR: PageRuler™ protein ladder, t20: soluble protein fraction after 20 h of expression, CFE: cell-free extract, FT: flow through, W: washing fraction, E1-E5: elution fractions (1 CV of elution buffer). NuPAGE™ Bis-Tris Mini Protein Gels, 4–12 %, Running conditions: 150 V for ~60 min in MES running buffer.

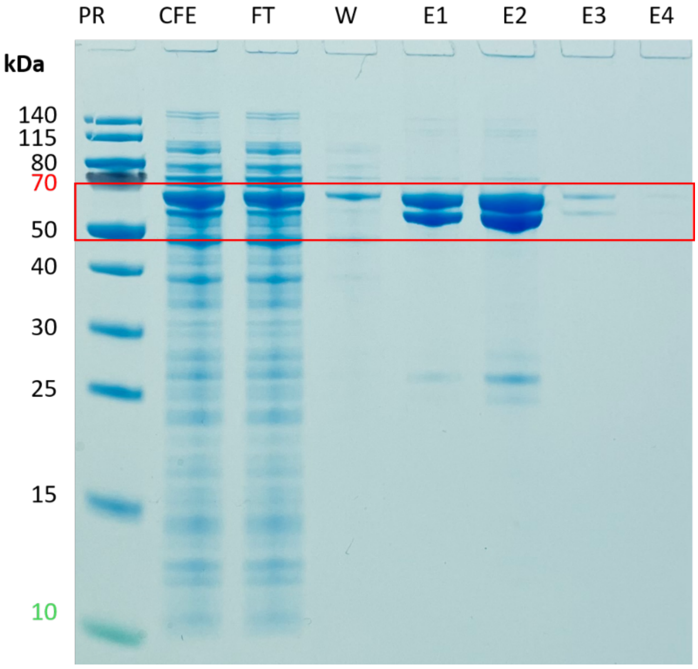

**Figure S4.** Stained SDS-PAGE showing protein fractions obtained during expression and the purification procedure of Oxy$_{\beta}$-G4S-Red fusion protein construct. The expected height for bands representing the large α-subunit (52.7 kDa) and the fused β-G4S-Red (58.4 kDa) are highlighted in red. PR: PageRuler™ protein ladder, CFE: cell-free extract, FT: flow through, W: washing fraction, E1-E4: elution fractions (1 CV of elution buffer). NuPAGE™ Bis-Tris Mini Protein Gels, 4–12 %, Running conditions: 150 V for ~60 min in MES running buffer.

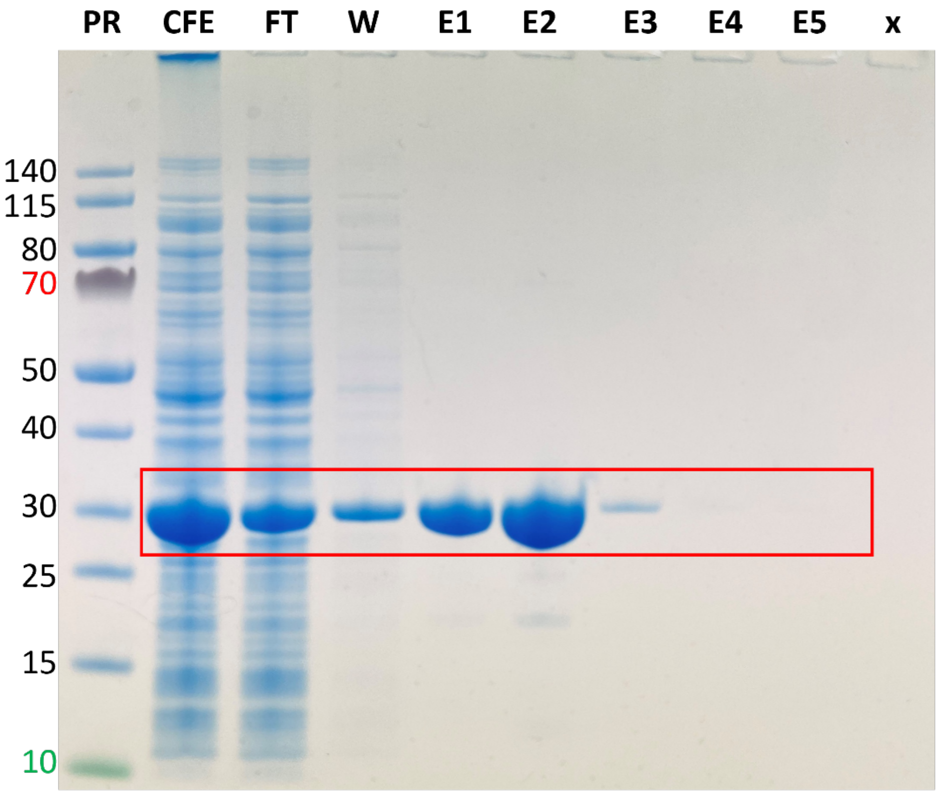

**Figure S5. Purification of GDH from *Bacillus megaterium* (29.0 kDa).** Stained SDS-PAGE showing protein fractions obtained during the purification procedure of GDH. The expected height for bands representing the desired protein is highlighted in red. PR: PageRuler™ protein ladder, CFE: cell-free extract, FT: flow through, W: washing fraction, E1-E5: elution fractions (1 CV of elution buffer). NuPAGE™ Bis-Tris Mini Protein Gels, 4–12 %, Running conditions: 150 V for ~60 min in MES running buffer.

### GC-FID Analysis

Parameters applied for non-chial GC-FID analysis are summarized in **Table S5.**

**Table S5. GC-FID parameters applied in this study**

| | |
|---|---|
| Gas chromatograph | Shimadzu GC-2010 Plus (Kyoto, Japan) |
| Column | Optima™ 5 MS GC from Macherey-Nagel™ |
| length<br>inner diameter<br>film thickness | 30 m<br>0.25 mm<br>0.25 µM |
| Injection volume | 1 µL |
| Injection temp. | 230 °C |
| Injection mode | Split |
| Carrier gas | $N_2$ |
| Flow control mode | Linear velocity |
| Pressure | 116.3 kPa |
| Total flow | 54.0 mL min$^{-1}$ |
| Column flow | 2.37 mL min$^{-1}$ |
| Linear velocity | 49.1 cm s$^{-1}$ |
| Purge flow | 3.0 mL min$^{-1}$ |
| Split ratio | 20.5 |
| Oven temp. program | 70 °C, 5 °C min$^{-1}$ to 110 °C, 15 °C min$^{-1}$ to 280 °C, hold for 5 min |
| FID temperature | 250 °C |
| Retention times | indene: 5.91 min<br>acetophenone (IS): 6.27 min<br>1*H*-indenol: 9.57 min<br>*cis*-indanediol: 12.50 min |

Product quantification via GC-FID was conducted by using the linear regression of recorded calibration curves (**Figures S6** and **S7**). In all measurements, acetophenone was used as an internal standard (IST).

#### Calibration of 1*H*-indenol

Within the period of this study, compound 1*H*-indenol was not commercially available and its synthesis was reported as challenging due to the formation of explosives.[3] Because of that, we used a calibration curve for 1-indanone (**Figure S6**), which was used to evaluate the concentration of 1*H*-indenol by approximation via the relative response factor (RRF).[1]

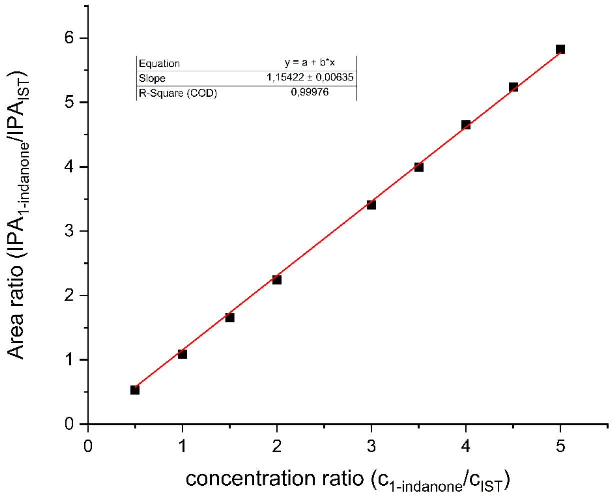

**Figure S6**. Calibration curve for 1-indanone using acetophenone as IST with non-chiral GC-FID. Applied GC parameters can be found in Table S5. The linear regression is highlighted in red.

#### Calibration of *cis*-indanediol

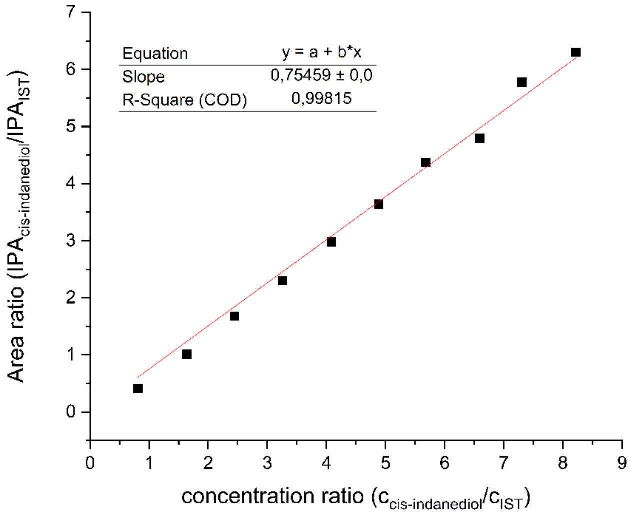

**Figure S7**. Calibration curve for cis-indanediol using acetophenone as IST with non-chrial GC-FID. Applied GC parameters can be found in Table S5. The linear regression is highlighted in red.

#### Example chromatograms obtained from GC-FID analyses

Example chromatograms obtained from measuring *in vitro* reactions highlighting the peaks of interest are depicted in **Figure S8**.

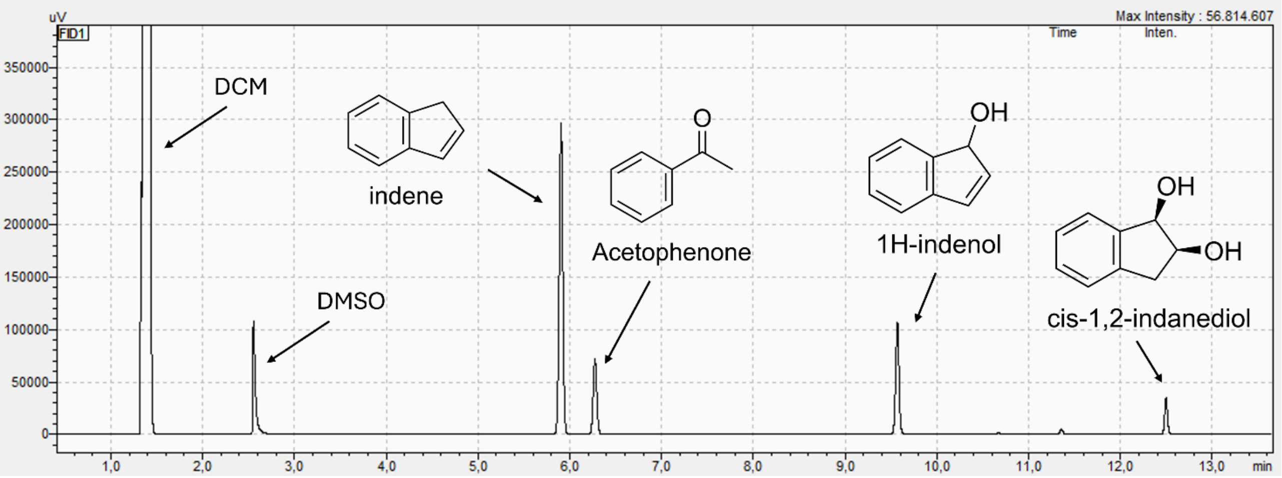

**Figure S8.** Extract from the chromatogram obtained from GC-FID analysis with parameters from **Table S5**. RT: 1.364 min dichloromethane (DCM), 2.554 min DMSO, 5.906 min indene, 6.274 min acetophenone, 9.571 min 1*H*-indenol, 12.497 min *cis*-1,2-indanediol.
